# Ventral Hippocampal Temporoammonic and Schaffer Collateral Pathways Differentially Control Fear- and Anxiety-Related Behaviors

**DOI:** 10.1101/2025.11.26.690696

**Authors:** Maltesh Kambali, Muxiao Wang, Rajasekar Nagarajan, Jinrui Lyu, Howard Gritton, Uwe Rudolph

**Author notes:** **Corresponding Author** Uwe Rudolph, Department of Comparative Biosciences, College of Veterinary Medicine, University of Illinois Urbana-Champaign, 3520 Veterinary Medicine Basic Science Building, 2001 S Lincoln Ave, Urbana, IL 61802-6178.

## Abstract

The ventral hippocampus plays a crucial role in regulating anxiety- and fear-related behaviors. Previously, we demonstrated that diazepam reduces anxiety-like behavior by inhibiting the dentate gyrus and CA3 principal neurons via α2-GABA_A_Rs, while inhibition of CA1 pyramidal neurons is necessary to suppress fear-related responses. This study investigated the role of inputs from ventral CA3 (vCA3) and entorhinal cortex to ventral CA1 (vCA1) in anxiety- and fear-like behavior using bidirectional optogenetic modulations. Adult C57BL/6J male and female mice were subjected to bilateral stereotaxic injection of a viral vector expressing channelrhodopsin or halorhodopsin into vCA3 or into layers II-III of lateral entorhinal cortex, followed by bilateral implantation of fiberoptic ferrules into vCA1. After four weeks of recovery, mice were assessed for anxiety-like behavior in the novel open field, elevated plus maze, and Vogel conflict tests, and by contextual and trace fear conditioning for fear. The behavior of the mice was recorded under laser ‘ON’ and ‘OFF’ conditions in all experiments. The activation of vCA3 to vCA1 projections (i.e., Schaffer collateral pathway) increased anxiety- and fear-related behaviors, whereas inhibition reduced such behaviors. In contrast, optogenetic activation or inhibition of EC to vCA1 projections (i.e., temporoammonic pathway) had no effect on anxiety-related behavior but positively or negatively modulated fear-related behavior, respectively. These results suggest that while fear-related behavior is modulated by both inputs to vCA1, modulation of anxiety-related behavior is input-specific for the vCA3 to vCA1 projection. In summary, this study offers mechanistic insights into the complex organization of hippocampal circuitry underlying fear and anxiety.

**GRAPHICAL ABSTRACT:** 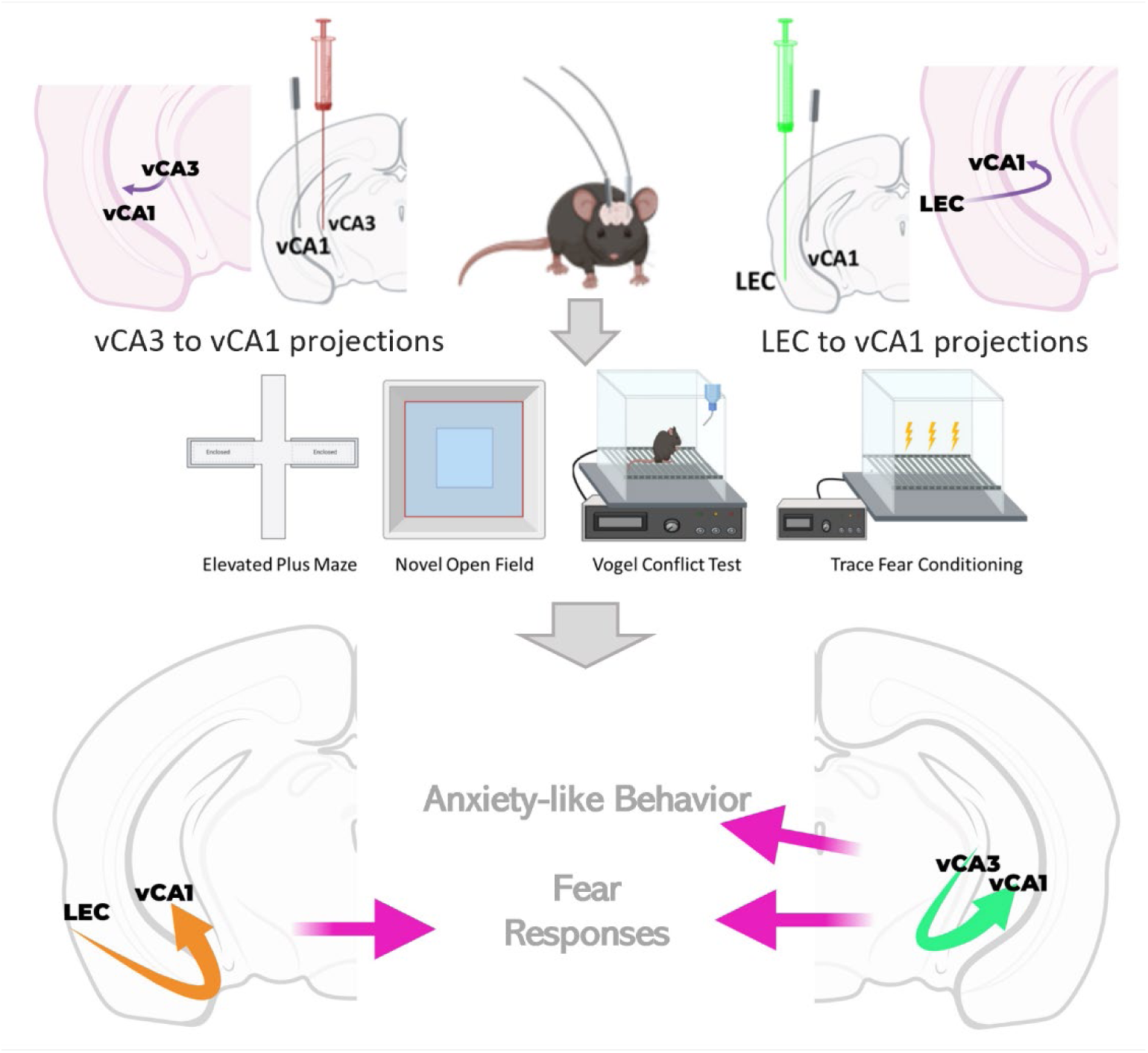

## 1. INTRODUCTION

Fear and anxiety are subjective experiences in humans. The differentiating factor is the origin (threat) which can be real or perceived and the extent of responses. They are negative valence states triggered by distinct environmental stimuli (such as vague, unidentified threats versus immediate, objectively danger situations), each leading to distinct defensive responses (heightened alertness and risk assessment versus freezing, fighting, or active avoidance) (1–3). In literature describing pre-clinical studies, the terms of fear and anxiety are used vaguely, and the distinction is questionable compared to observations in humans. A simplified convention for studies aiming to identify neuronal circuits with a minimal distinction based on the type of behavior evaluated in rodents is to use the terms fear-related and anxiety-like behaviors.

Decades of research have identified the role of different brain areas in the circuitry of fear and anxiety(1–3). However, until recently, existing methodologies have lacked the spatial and temporal precision needed to discern specific cell types, projections, and their roles within broader neural networks. Recent investigations focusing on neural circuits have shown that, although there is some convergence in the neurocircuitry and neuronal populations involved in the regulation of fear and anxiety(4,5), there are also notable differences(6,7). These findings highlight the complexity and specialization within the circuits that mediate these closely related emotional states. Neuroimaging studies have implicated hippocampal dysfunction in mood and anxiety disorders(6,8–10). In line with its established roles in cognitive and emotional processes, the hippocampus demonstrates significant variation along its dorsoventral axis, particularly in relation to its afferent and efferent functional connectivity. This gradient of connectivity reflects the differential involvement of dorsal regions in spatial memory and cognitive functions, while ventral regions are more tightly associated with emotional regulation and affective processing(6,11,12).

For cognitive hippocampal functions, distinct roles of the trisynaptic and temporoammonic (TA) pathways have been identified(13,14)However, this is not the case for emotional hippocampal functions. Circuit-level approaches to study emotional regulation in the hippocampus have been limited so far. The hippocampal circuitry includes predominantly unidirectional excitatory projections from the entorhinal cortex to DG to CA3 to CA1 (trisynaptic pathway), and direct projections from the entorhinal cortex to CA1 (temporoammonic pathway). However, the role of hippocampal circuitry in fear and anxiety remains largely unexplored.

Recent insights into the role of ventral CA1 (vCA1) projections in the modulation of fear and anxiety reveal that vCA1 projections to other regions, such as the amygdala, are necessary for contextual fear memory(15), and projections to hypothalamus are required for anxiety-like behavior(16). While optogenetic activation of Hoxb8-positive microglial cells in vCA1 increases anxiety-like behavior(17), the synchronized activity of vCA1 and medial prefrontal cortex is associated with anxiety(18). Previously, we explored the role of hippocampal subregions in anxiety- and fear-related behavior utilizing novel cell type- and region-specific conditional knockouts of the GABA_A_ receptor α2 subunit, thereby revealing subregion-specific mechanisms through which benzodiazepines exert their behavioral effects. We discovered that α2-containing GABA_A_ receptors in CA3 and dentate gyrus (DG) pyramidal neurons are required for reduction of anxiety-like behavior but not of fear-related behavior by the benzodiazepine diazepam, indicating that modulation of excitatory microcircuits within these subregions critically gates anxiety-like behavior. Conversely, α2-containing GABA_A_ receptors in CA1 pyramidal neurons were selectively necessary for diazepam’s reduction of fear-related behavior, but their deletion in these neurons did not alter the drug’s anxiolytic effects. Together, these findings demonstrate a functional segregation in intrahippocampal circuits, wherein diazepam engages α2-GABA_A_Rs in DG and CA3 to modulate anxiety, while recruiting α2-GABA_A_Rs in CA1 to modulate fear. This work provides mechanistic insight into how benzodiazepines achieve distinct behavioral actions through differential engagement of inhibitory signaling within hippocampal subregions, highlighting that expression of fear-related behaviors involves complex excitatory inhibitory computation in CA1, which apparently can be overridden by the CA3 inputs in a context-dependent manner to exhibit anxiety-like behaviors. This suggests that the computation in CA1 may depend on an input (e.g., potentially a direct entorhinal cortical input) determining how CA1 integrates incoming information and enabling the expression of distinct behavioral outcomes according to the context. This study thus demonstrated a double dissociation in the regulation of anxiety-like versus fear-related behaviors at the level of intra-hippocampal circuits(19). This insight into the differential involvement of hippocampal subregions led us to examine the role of projections to CA1 in the regulation of anxiety-like and fear-related behaviors. Based on the aforementioned pharmacological studies we employed bidirectional optogenetic modulation of ventral hippocampal monosynaptic excitatory projections to test the hypothesis that the CA3 to CA1 Schaffer collateral projections are essential to modulate anxiety-like behavior, while the direct temporoammonic pathway projections from EC to CA1 would modulate fear-related behavior.

## 2. RESULTS

### 2.1 Investigation of Projections from Ventral CA3 Pyramidal Neurons to Ventral CA1

We set out to investigate the role of vCA3-vCA1 and EC-vCA1 projections for the modulation of anxiety-like behavior by optogenetic manipulation of their activity. To this end, an AAV expressing either channelrhodopsin or halorhodopsin was stereotaxically injected into EC or vCA3, and a fiberoptic ferrule was implanted into vCA1 (**Fig. 1A and Fig. 3A**). Four weeks later, mice were subjected to behavioral paradigms assessing anxiety- or fear-related behavior, specifically the elevated plus maze, novel open field, Vogel conflict test, and trace fear conditioning paradigms (**Fig. 1B and Fig. 3B**). C-fos staining revealed that the fiberoptic illuminations of projections from vCA3 principal neuronal projections to vCA1 led to neuronal activation (channelrhodopsin) or inhibition (halorhodopsin), respectively based on percent c-fos positive cell counts **(Fig. 1D and E**). One-way ANOVA [F (5,42) = 83.93, p<0.0001] followed by Sidak’s multiple comparison test showed that the optogenetic activation of vCA3 projections to vCA1 led to an increase in c-fos positive cells (t=15.07, p<0.0001) and the optogenetic inhibition of vCA3 projections to vCA1 led to a decrease in c-fos positive cells (t=3.122, p=0.0161) compared to the sham control. There was no significant change in c-fos positive cells in the dorsal CA1 (**Fig. 1D)**.

**Figure 1.**
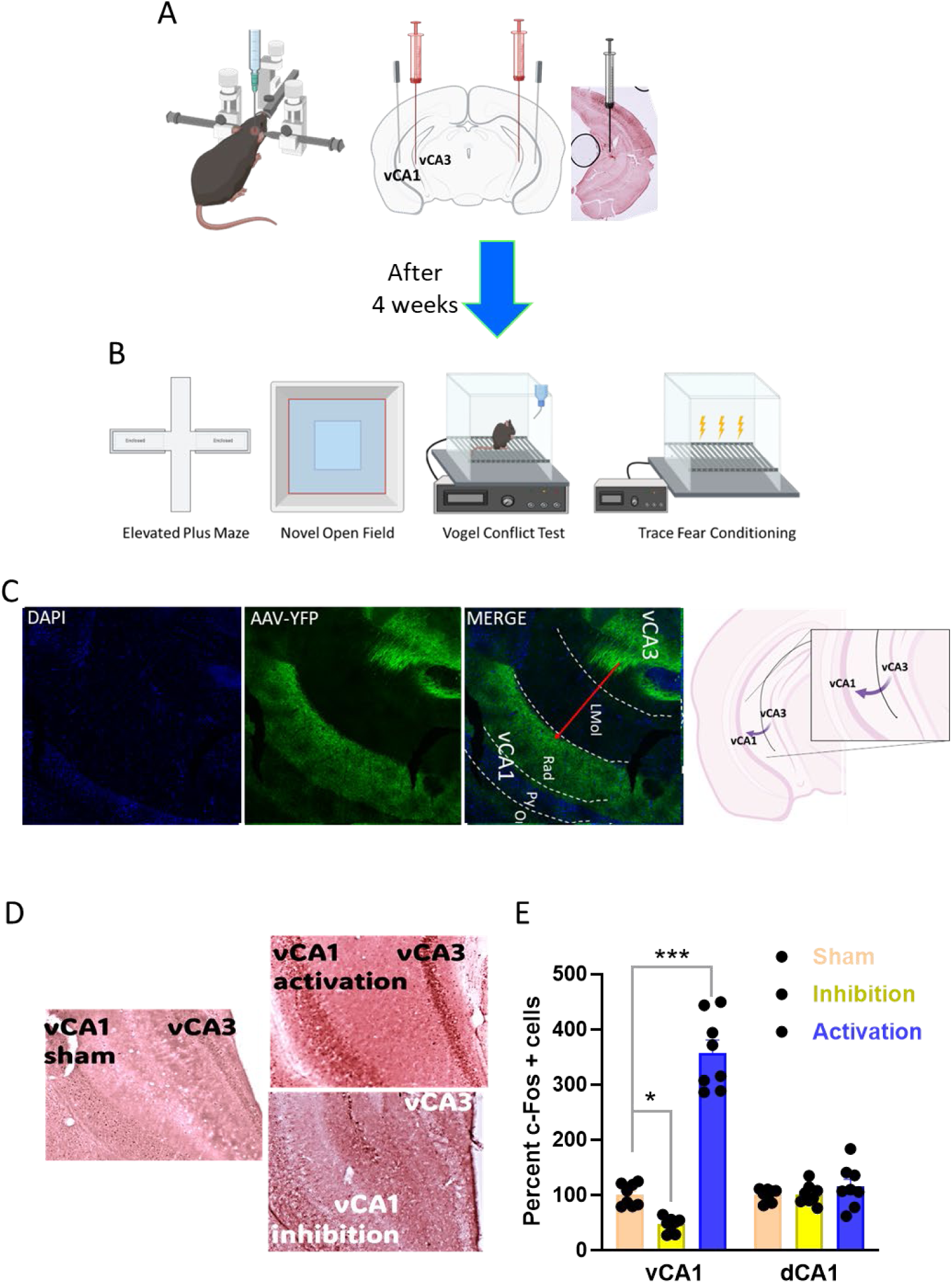
Optogenetic inhibition or activation of vCA3-vCA1 projections with halorhodopsin (eNpHR3.0) or channelrhodopsin [hChR2(H134R)]. A. Schematics of stereotaxic bilateral injection of viral vectors into vCA3 and implantation of the fiberoptic ferrules into vCA1, including images of viral injection site, c-fos staining (coronal section). B. Schematics of behavioral analysis. C. Opsin viral vector immunofluorescence. The immunofluorescence images showing the fluorescently labelled viral injection site in vCA3, and the projections to vCA1 are largely terminating in the radial region (Rad) of vCA1, i.e., proximal to the vCA1 pyramidal cell bodies (Py). Last image is schematics of vCA3 projections to vCA1. D. c-fos staining images showing the neuronal activation (upper image) or inhibition (lower image) in vCA1 by optogenetic illumination of vCA3 projections. E. The level of c-fos positive cells in ventral and dorsal CA1 with illumination of channelrhodopsin (activation) or halorhodopsin (inhibition). n=8 each group. Data are presented as means ± SEM.

### 2.2 Neuronal Projections from Ventral CA3 Principal Neurons to CA1 Modulate Anxiety-Like Behavior

To visualize the sites of opsin viral vector injection in the vCA3 region of the hippocampus, we mounted YFP-expressing tissue on DAPI covered slides (**Fig. 1C**). In **Fig. 1C**, the merged image shows that the viral vector’s yellow fluorescence contrasted against a blue nuclear stain background. The merged image also indicates that the vCA3 collateral axonal projections synapse onto vCA1 dendrites in the stratum radiatum i.e., proximal to the vCA1 pyramidal cell layer bodies.

We first assessed anxiety-like behavior in the elevated plus maze. Optogenetic activation of vCA3 principal neuronal projections to vCA1 (ON) led to a decreased open arm time (**Fig. 2C**) compared to the OFF sessions in same animals. In contrast, optogenetic inhibition of the vCA3 principal neuronal projections to vCA1 (ON) increased the open arm time in the elevated plus maze **(Fig. 2C**). One way ANOVA [F (5, 47) = 4.405, P=0.0023] followed by Sidak’s multiple comparisons test showed significantly increased open arm exploration when vCA3 projections are inhibited in vCA1 (yellow laser ON, t=3.200, p=0.0049) and a significantly decreased open arm exploration when vCA3 projections are activated in vCA1 (blue laser ON, t=2.434, p=0.0372). The total distance travelled was not significantly different after optogenetic activation or inhibition (**Fig. 2D**). These results of activation and inhibition of vCA3 projections to vCA1 indicate that the vCA3 principal neuronal projections to vCA1 bidirectionally modulate anxiety-like behavior.

**Figure 2.**
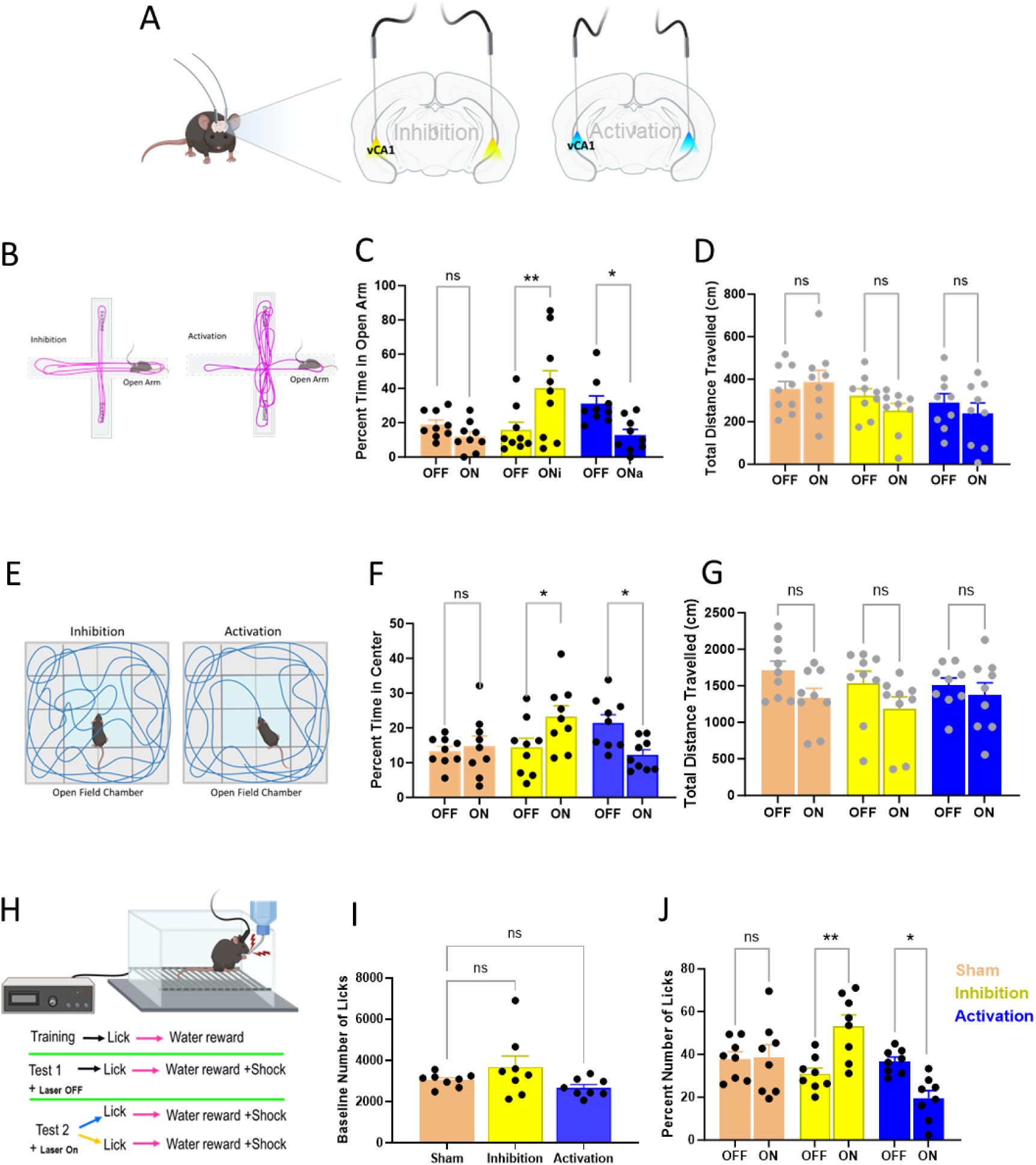
Illumination of vCA3 pyramidal neuronal projections at vCA1 modulates anxiety-like behavior. A. Coronal schematics of the brain showing fiber optic laser illumination site at ventral CA1 with blue (activation) or yellow (inhibition) lasers. B.-D. An elevated plus maze (schematic in B.) was performed to assess anxiety-like behavior coupled with optogenetic laser illumination. The bar graph shows that activation or inhibition of vCA3 neuronal projections to vCA1 decreased or increased the open arm exploration time the in elevated plus maze, respectively (C), while the total distance travelled in the elevated plus maze was not altered by any optogenetic illuminations (D). E.-G. A novel open field exploration test (schematic in E.) was performed to assess anxiety-like behavior with optogenetic laser stimulation. The bar graph shows that the activation or inhibition of vCA3 neuronal projections to vCA1 decreased or increased the center exploration in novel open field test, respectively (F). The total distance travelled in the open field test was not altered by optogenetic activation or inhibition (G). H.-J. A Vogel conflict test (schematic in H.) was performed to assess anxiety-like behavior in an approach-avoidance conflict. The baseline (without shock) number of licks was similar in all groups (I). Vogel task coupled with optogenetic illuminations over the test period to assess anxiety-like behavior. The bar graph shows that activation or inhibition of vCA3 projections to vCA1 decreased or increased the percent number of licks performed by the mice, respectively (J). n=8-9 (4-5 males and 4 females). Data are presented as means ± SEM.

In the novel open field, another paradigm used to assess anxiety-like behavior, optogenetic activation of vCA3 principal neuronal projections to vCA1 decreased the time spent in the center of the open field chamber (**Fig. 2F**) compared to the OFF session in same animals. In contrast, optogenetic inhibition of the vCA3 principal neuronal projections in vCA1 led to an increase in the time spent in the center of the open field chamber (**Fig. 2F**) compared to the OFF sessions.

A one way ANOVA [F (5, 48) = 3.564, P=0.0081] followed by Sidak’s multiple comparisons test showed significantly increased center exploration in the open field test when vCA3 projections are inhibited in vCA1 (yellow laser ON, t=2.651, p=0.0325), and a significantly decreased center exploration in the open field test when vCA3 projections are activated in vCA1 (blue laser ON, t=2.596, p=0.0375). The total distance travelled was not changed by optogenetic activation or inactivation (**Fig. 2G**), indicating that the locomotor activity in the novel open field was unaltered by optogenetic activation or inhibition. These results from two different anxiety-like behavior assessment paradigms, (i.e., elevated plus maze and novel open field), suggest that vCA1 mediates anxiety-like behavior and that its activity is modulated by inputs from vCA3 principal neuronal projections. In summary, vCA3 projections to vCA1 specifically regulate activity of vCA1 neurons, bidirectionally to modulate anxiety-like behavior.

Subsequently, to test another aspect of anxiety-like behavior, mice were subjected to the Vogel conflict test, which is based on an approach-avoidance conflict(21) Water-deprived mice are allowed to drink from a spout that delivers an electrical shock to the tongue, creating a conflict between the desire to drink and the desire to avoid a noxious stimulus (**Fig. 2H**). It has been developed to screen compounds for anxiolytic-like properties. A reduction of spout licking compared to the no-shock condition may be a component of anxiety-like behavior, while an increase reflects anxiolytic-like behavior(21–23). We developed a protocol in which animals undergo two counterbalanced test sessions 24 hours apart: once with laser ON and once with laser OFF. The baseline number of licks on the third day pre-test (i.e., without shock) was similar in all the groups tested (**Fig. 2I**). Optogenetic activation of vCA3 principal neuronal projections to vCA1 (ON test sessions) led to a decrease in the percent number of licks performed compared to the no illumination (i.e., OFF test session) in the same animals (**Fig. 2J**). In contrast, the optogenetic inhibition of the vCA3 principal neuronal projections to vCA1 increased the number of licks performed (**Fig. 2J**). A one-way ANOVA [F (5, 42) = 7.154, P<0.0001] followed by Sidak’s multiple comparisons test showed significantly increased number of licks when vCA3 projections are inhibited in vCA1 (t=3.831, p=0.0013), and a significantly decreased number of licks when vCA3 projections are activated in vCA1 (t=2.977, p=0.0144). The number of licks without shock during the pretest (Baseline) was consistent across all experimental groups (**Fig. 2I**). However, this number decreased to approximately 40-50% during the test session following the introduction of a shock, which induced a conflict-avoidance response. The shock-induced reduction in licking behavior was modulated bidirectionally through optogenetic manipulations of vCA3 projections to vCA1.These results indicate that anxiety-like behavior in this conflict-avoidance approach is modulated by the vCA3 principal neuronal projections to vCA1 bidirectionally.

### 2.3 Projections from Ventral CA3 Principal Neurons to Ventral CA1 Modulate Fear-Related Behavior

In order to evaluate the role of the vCA3 neuronal projections to vCA1 for fear expression, we performed contextual and trace fear conditioning (**Fig. 3A**), evaluating fear expression related to context (**Fig. 3B**) and to tone (**Fig. 3C**) separately. The lasers were OFF during the conditioning session and ON during the two tests sessions (for context, and for tone). Optogenetic activation of vCA3 neuronal projections to vCA1 increased the freezing in the same context that was used for fear conditioning, compared to the sham control animals (**Fig. 3B**). In contrast, optogenetic inhibition of the vCA3 principal neuronal projections to vCA1 decreased the freezing behavior in the context that was used for fear conditioning (**Fig. 3B**). One way ANOVA [F (2, 24) = 21.63, P<0.0001] followed by Sidak’s multiple comparisons test showed significantly increased fear expression when vCA3 projections are activated in vCA1, (t=4.021, p=0.00010) and that the optogenetic inhibition of vCA3 projections to vCA1 significantly decreased freezing (t=2.497, p=0.0365). These results indicate that vCA3 principal neuronal projections to vCA1 bidirectionally modulate contextual fear expression.

**Figure 3.**
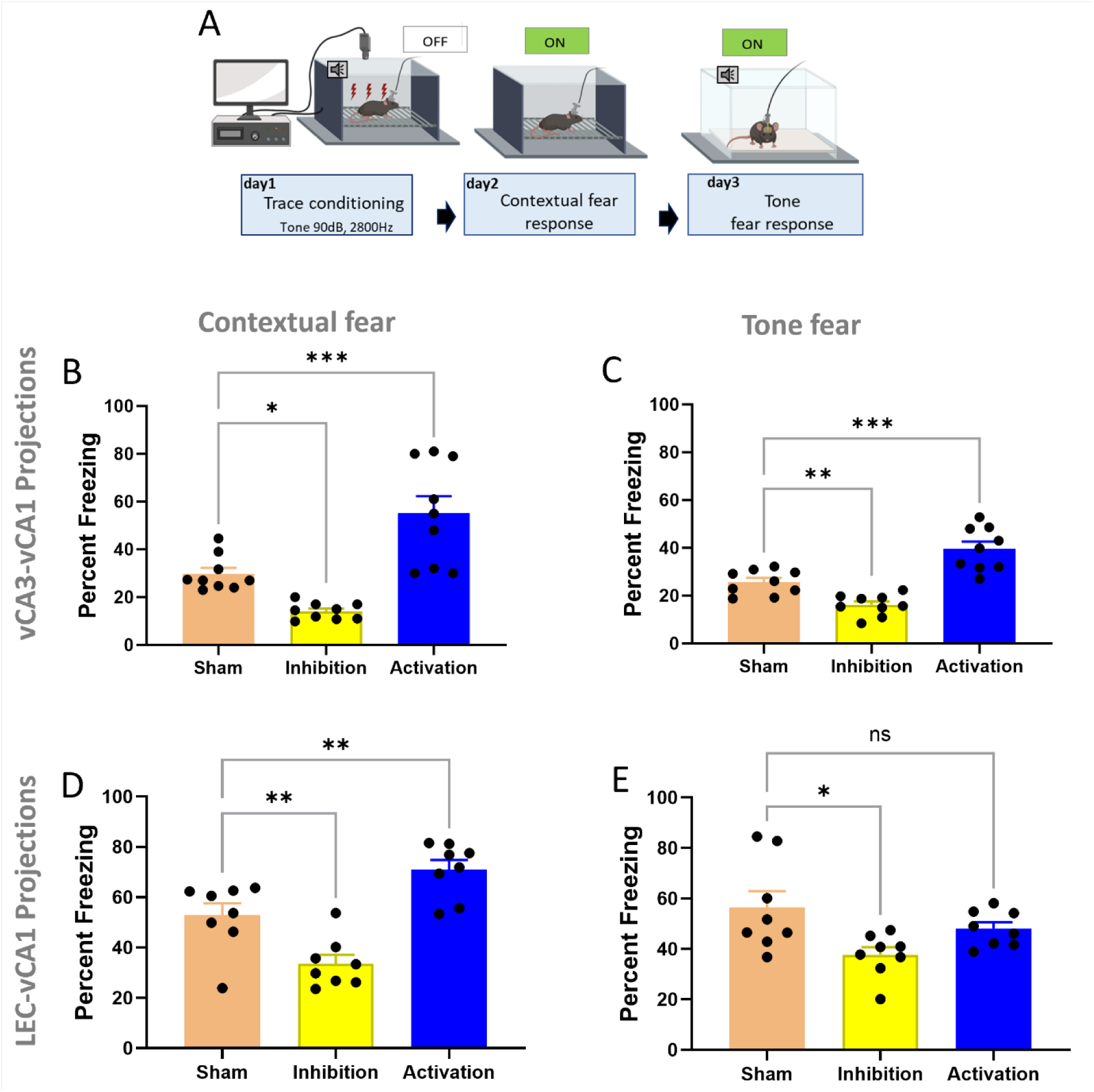
Projections from both vCA3 and lateral entorhinal cortex to vCA1 modulate fear memory retrieval. A. Schematics show the trace fear conditioning procedure for optogenetic modulation of vCA3 or LEC projections at vCA1 during the context- and tone-related fear expression. B., D. Contextual fear expression testing in the same chamber that was used for conditioning. Activation or inhibition of vCA3 projections (B) or LEC projections (D) to vCA1 Increased or decreased, respectively, the freezing behavior or fear expression to context. C., E. Fear expression in response to the tone in a new context. Activation or inhibition of vCA3 projections to vCA1 increased or decreased, respectively, the freezing behavior or fear expression in response to the tone (C), while inhibition of the projections from LEC to vCA1 decreased the freezing behavior or fear expression in response to the tone (E). n=8-9 (4-5 males and 4 females). Data are presented as mean ± SEM.

Further, the fear expression to the conditioned tone test showed that the optogenetic activation of vCA3 principal neuronal projections to vCA1 increased the freezing to the tone compared to the sham control animals (**Fig. 3C**). In contrast, optogenetic inhibition of the vCA3 principal neuronal projections to vCA1 decreased the freezing behavior in response to the tone (**Fig. 3C)**. One-way ANOVA [F (2,21) = 4.726, p=0.0202] followed by Sidak’s multiple comparison test showed that the optogenetic activation of vCA3 projections to vCA1 increases freezing (t=4.515, p=0.0003) and that the optogenetic inhibition of vCA3 projections in vCA1 significantly decreased freezing (t=3.096, p=0.0098). These results indicate that vCA3 principal neuronal projections to vCA1 bidirectionally modulate fear expression following trace fear conditioning.

In summary, these results suggest that – in addition to the modulation of anxiety-like behavior - the intrahippocampal projections from vCA3 neurons to vCA1 are involved in the modulation of fear-related behaviors (**Supplementary Fig. 1**).

### 2.4 Investigation of Entorhinal Cortical Projections to Ventral CA17

To explore the role of the direct projection from EC to CA1 we used a similar approach to study anxiety- and fear-related behaviors. The immunofluorescence images show the target location of opsin viral vector injection into the LEC (**Fig. 4A and C**) in the second row of Fig. 3C indicates that the LEC principal neuronal projections largely synapse onto vCA1 dendrites in the molecular later (LMol) i.e., distal to the vCA1 pyramidal cell layer bodies(24,25). The quantification of c-fos positive cells revealed that the fiberoptic illuminations of LEC principal neuronal projections to vCA1 led to neuronal activation (channelrhodopsin) or inhibition (halorhodopsin), respectively (**Fig. 4D and E**). A one-way ANOVA [F (5,42) = 37.87, p<0.0001] followed by Sidak’s multiple comparison test shows that the optogenetic activation of LEC principal neuronal projections in vCA1 led to increase in c-fos positive cells (t=8.174, p<0.0001) and the optogenetic inhibition of LEC principal neuronal projections in vCA1 led to decrease in c-fos positive cells (t=5.441, p<0.0001) compared to sham control. There was no significant change in c-fos positive cells in the dorsal CA1.

**Figure 4.**
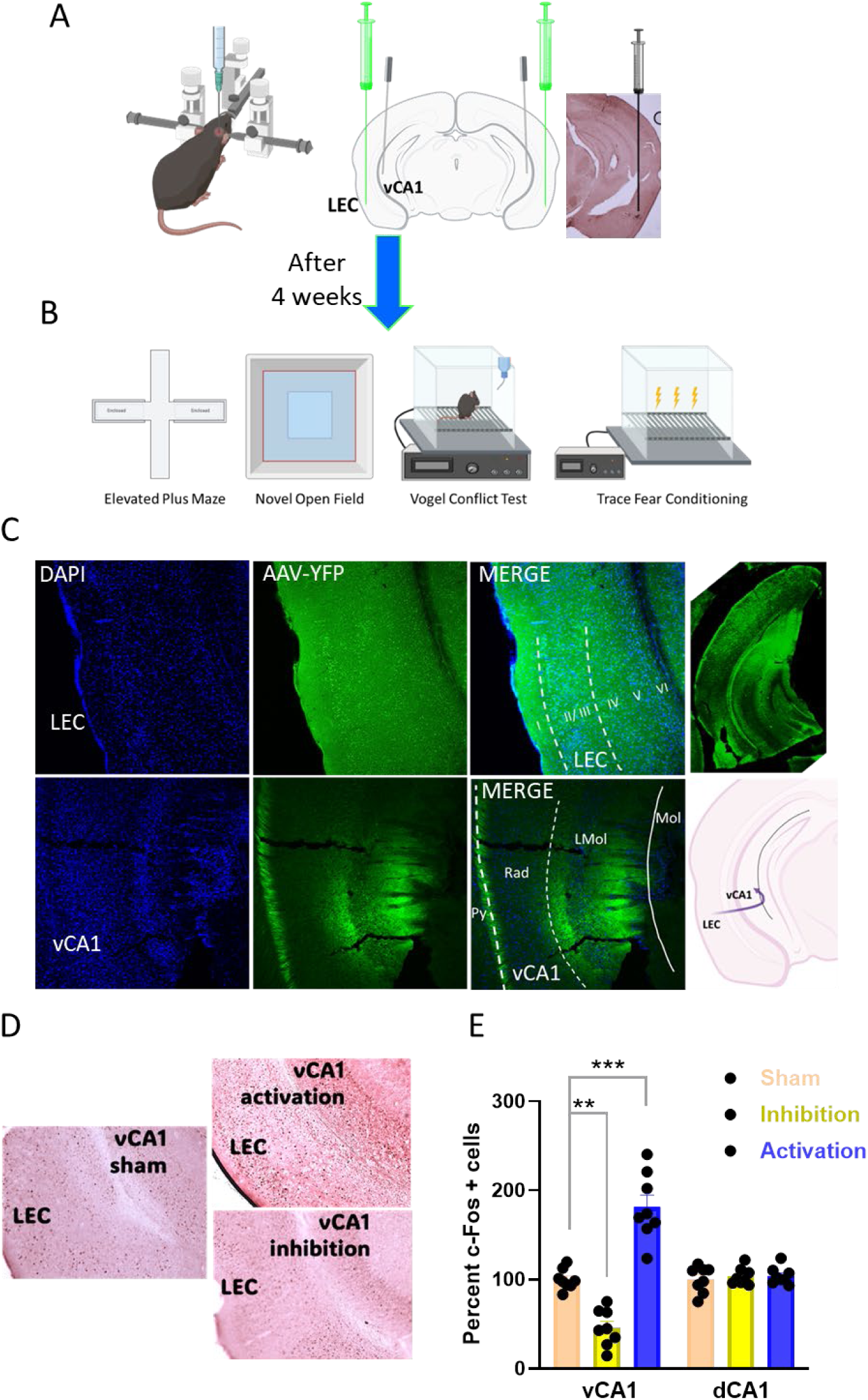
Optogenetic inhibition or activation of lateral entorhinal cortex (LEC) to vCA1 projections with halorhodopsin (eNpHR3.0) or channelrhodopsin [hChR2(H134R)]. A. Schematics of bilateral injection of viral vectors into LEC, and implantation of the fiber optic ferrule into vCA1, including images of viral injection site- c-fos staining (coronal section). B. The behavioral experiments performed after surgical recovery include anxiety-like behaviors (elevated plus maze, novel open field, Vogel conflict test) and trace fear conditioning for fear expression. C. Top row: immunofluorescence images showing the opsin viral vector fluorescence in layer 2/3 LEC region. Bottom row: LEC projections to vCA1 are largely terminating in the lacunosum molecular region (LMol) of vCA1, i.e., distal to the vCA1 pyramidal cell bodies (Py). The right image in the top row is showing the target opsin viral injection site, and the right image in the lower row the schematics of LEC projections to vCA1. D. c-fos staining images showing neuronal activation (top image) or inhibition (bottom image) in vCA1 by optogenetic illumination of LEC projections. E. The level of c-fos positive cells in ventral and dorsal CA1 with illumination of channelrhodopsin (activation) or halorhodopsin (inhibition). n=8 (4 males and 4 females). Data are presented as means ± SEM.

### 2.5 Lateral Entorhinal Cortical Projections to Ventral CA1 do not Modulate Anxiety-Like Behavior

Optogenetic activation or inhibition of LEC principal neuronal projections to vCA1 (ON) did not change the open arm time in the elevated plus maze compared to the OFF session in same animals (**Fig. 5C**). One-way ANOVA [F (5,45) = 3.738, p=0.0065] followed by Sidak’s multiple comparison test showed that the optogenetic activation of LEC principal neuronal projections to vCA1 does not alter the time in the open arms (t=1.123, p=0.6071) and that the optogenetic inhibition of LEC principal neuronal projections to vCA1 does not alter the time in the open arms (t=0.4180, p=0.9666). Moreover, there was no change in the total distance travelled (**Fig. 5D)**. These results indicate that LEC principal neuronal projections to vCA1 neurons do not modulate anxiety-like behavior in the elevated plus maze.

**Figure 5.**
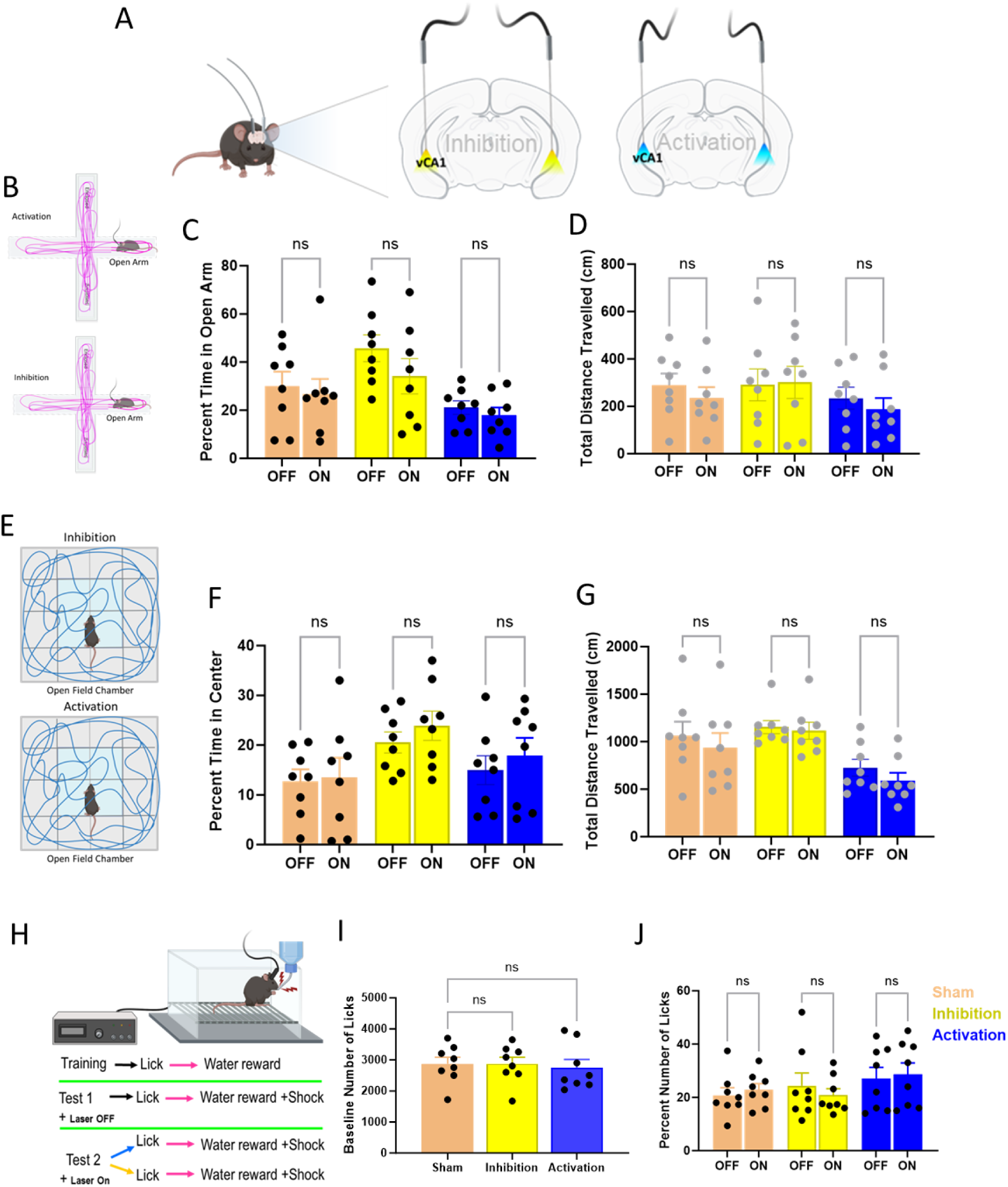
Illumination of lateral entorhinal cortical projections in vCA1 modulates anxiety-like behavior. A. Coronal schematics of the brain showing the fiber optic laser illumination site at ventral CA1 with blue (activation) or yellow (inhibition) lasers. B.-D. An Elevated plus maze (schematic in B.) was performed to assess the anxiety-like behavior coupled with optogenetic laser illumination. C. The bar graph shows that activation or inhibition of LEC projections to vCA1 does not alter the open arm exploration in elevated plus maze and the total distance travelled (D.) in the elevated plus maze. E. Novel open field exploration test performed to assess anxiety-like behavior with optogenetic laser illumination. F. The bar graph shows that the activation or inhibition of LEC projections to vCA1 does not alter the center exploration and the G. total distance travelled in novel open field test. H. Vogel conflict test in the schematics showing the experimental protocol in the optogenetic illumination coupled Vogel task. I. Vogel conflict test with baseline number of licks (without shock) was similar in all groups. J. Vogel task coupled with optogenetic stimulations over the test period to assess anxiety-like behavior. The bar graph showing that activation or inhibition of vCA3 projections at vCA1 does not alter the percent number of licks performed by the mice, respectively. n=8 (4 males and 4 females). Data are presented as means ± SEM.

In the novel open field, optogenetic activation or inhibition of LEC principal neuronal projections to vCA1 (ON) did not alter the time spent in the center of the open field chamber compared to the OFF sessions in same animals (**Fig. 5F**). One-way ANOVA [F (5,45) = 1.465, p=0.2201] followed by Sidak’s multiple comparison test showed that the optogenetic activation of LEC principal neuronal projections to vCA1 does not alter the time in center of the open field (t=0.3701, p=0.9764) and that the optogenetic inhibition of LEC principal neuronal projections to vCA1 does not alter the time in the center of the open field (t=0.6302, p=0.8973). Moreover, there was no change in the total distance travelled in the open field (**Fig. 5G**). These results suggest that LEC principal neuronal projections to vCA1 have no role in modulation of anxiety-like behavior in the open field.

Optogenetic activation or inhibition of LEC principal neuronal projections to vCA1 (ON) do not alter the percent number of licks performed compared to the no illumination (i.e., ‘OFF’) test session in the same animals (**Fig. 5J**). One-way ANOVA [F (5,44) = 0.7351, p=0.6011] followed by Sidak’s multiple comparison test showed that the optogenetic activation of LEC principal neuronal projections to vCA1 had no effect on spout licking (t=0.0749, p=0.9998) and that the optogenetic inhibition of LEC principal neuronal projections to vCA1 had no effect on spout licking (t=0.2893, p=0.9884). There was no change in the baseline number of licks (without shock) performed (**Fig. 5I**). These results indicate that LEC principal neuronal projections to vCA1 do not modulate conflict-avoidance behavior in the Vogel conflict test.

### 2.6 Lateral Entorhinal Cortical Projections to Ventral CA1 Modulate Fear-Related Behavior

To evaluate the role of LEC principal neuronal projections to vCA1 on fear expression, we performed a trace fear conditioning paradigm as described above and in **Fig. 3A**. In the contextual fear expression test optogenetic activation of LEC principal neuronal projections to vCA1 increased the freezing behavior compared to the sham control animals (**Fig. 3D**). In contrast, the optogenetic inhibition of the LEC principal neuronal projections to vCA1 decreased the freezing behavior in same context that was used for fear conditioning (**Fig. 3D**). One way ANOVA [F (2, 22) = 24.09, P<0.0001] followed by Bonferroni’s multiple comparisons test showed significantly increased fear expression when LEC projections are activated in vCA1 (t=3.633, p=0.0029), and a significantly decreased fear expression when LEC projections are inhibited in vCA1 (t=3.203, p=0.0082) These results indicate that LEC principal neuronal projections to vCA1 bidirectionally modulate contextual fear expression. There is a different level of freezing response in sham controls for both projections, i.e., vCA3 to vCA1 and LEC to vCA1, as the experiments were not performed at the same time.

Further, the fear expression test in response to the tone showed that the optogenetic activation of LEC principal neuronal projections to vCA1 did not affect the freezing behavior to the tone (**Fig. 3E**). In contrast, optogenetic inhibition of the LEC principal neuronal projections to vCA1 decreased the freezing behavior to the tone (**Fig. 3E**). One way ANOVA [F (2, 22) = 5.425, P=0.0122] followed by Sidak’s multiple comparisons test showed significantly decreased fear expression when LEC projections to vCA1 are inhibited (t=3.277, p=0.0069) and the optogenetic activation of LEC principal neuronal projections to vCA1 do not alter the freezing (t=1.395, p=0.3225). These results indicate that LEC principal neuronal projections to vCA1 neurons bidirectionally modulate contextual fear expression following trace fear conditioning.

In summary, these results indicate that the projections from lateral entorhinal cortex to vCA1, while not modulating anxiety-like behaviors, are involved in the modulation of vCA1 principal neuronal activity related to contextual fear expression (**Supplementary Fig. 1**).

## 3. DISCUSSION

While a robust history of lesion studies and pharmacological/stimulation studies have established a role for the ventral hippocampus in fear-related and anxiety-like behaviors (12,16,26), the specific role of hippocampal circuit projections remains largely unknown. Learning processes require precise neuronal communication across distributed brain networks. Notably, very few studies have specifically explored the differential functions of the subregions within the ventral hippocampus in fear and anxiety. Jimenez et al. identified that the vCA1 neurons projecting to lateral hypothalamus (LH) mediate anxiety-like behavior(16) but CA1 neurons projecting to amygdala mediate contextual fear-related behaviors (18), suggesting that the vCA1 output relies on the computation of information which partly depends on the strengths of inputs to vCA1 from vCA3 and LEC and is mostly dependent on the context. Another study showed that the synchronized activity of vCA1 and medial prefrontal cortex is associated with anxiety-like behavior(18). In addition, Graham et al. showed that high frequency stimulation of vCA1 neurons reduces amygdala activity and inhibits fear responses(27). These studies implicate that vCA1 projections to different brain regions may be functionally distinct due to differential refinement of information depending on the complexity of sensory inputs with relevance for fear-related and anxiety-like behaviors. However, the roles of the hippocampal projections that modulate fear-related and anxiety-related behaviors are not yet well understood. Previously, our own work has shed some light on this aspect, showing that inhibition of dentate gyrus and CA3 principal neurons is necessary for the reduction of anxiety-like behavior by diazepam, as this response to diazepam is absent in mice lacking α2-containing GABA_A_ receptors (α2-GABA_A_Rs) in DG or CA3 principal neurons, respectively, whereas inhibition of CA1 principal neurons via α2-GABA_A_Rs is necessary for diazepam-induced suppression of fear responses, as this effect of diazepam is absent in mice lacking α2-containing GABA_A_ receptors (α2-GABA_A_Rs) in CA1 principal neurons(19). Taken together, this indicates that the hippocampal subregions may have different roles in modulating fear-related and anxiety-like behaviors. This study provided evidence that anxiety-like behaviors are modulated by the discrete neuronal cell type in defined regions of the hippocampal circuitry, however, it did not address the point whether inputs into CA1 from different sources (e.g., entorhinal cortex and CA3) would be specific for modulation of anxiety-like or fear-related behaviors. The present study was designed to address whether each of these pathways have specific contributions to modulating fear and anxiety directly. We found that both LEC to vCA1 and vCA3 to vCA1 projections modulate fear-related behaviors but differ in their ability to influence anxiety-like behaviors. In particular, only the vCA3 to CA1 projection was found to modulate both behaviors. These results are consistent with the view that the computation in vCA1 involves bundling the inputs from these two projections. They also indicate that depending on the context, one or both projections either synchronize or cancel out their influence on vCA1, thereby modulating its output. These findings are in line with a previous study showing that a colchicine-induced lesion of ventral dentate gyrus decreases the anxiety-like behavior in elevated plus maze and in open field tests(28) while another recent study showed that a selective knockdown of the GABA_A_ receptor α2 subunit in the dorsal dentate gyrus induces anxiety-like behavior(29). In contrast, Kheirbek et al. reported that activation of ventral dentate gyrus (DG) granule cells reduces anxiety-like behavior, but these neurons do not play a role in the retrieval of contextual fear memory(6). In contrast, we found that activating vCA3 projections to vCA1 increases anxiety-like behavior and enhances both contextual and tone fear memories. A separate study by Graham et al. reported that high-frequency stimulation of the vCA1 (vCA1) reduces amygdala activity and suppresses fear responses(27). Both of these studies(6,27) present results that differ from ours, and they also show discrepancies between each other, possibly due to differences in the stimulation protocols, e.g., the high-frequency stimulation may mimic a stressful situation. In our study, we noticed a remarkable similarities but also differences with the results from our previous work on conditional knockouts of the GABA_A_ receptor α2 subunit in CA1, CA3 and DG principal neurons. Whereas in the Engin et al. (2016) study the results obtained with the Vogel conflict test were in line with those obtained in a fear-potentiated startle test, but not with the elevated plus maze, light/dark choice, and reticular-elicited theta tests, suggesting that it detects fear-related behavior rather than anxiety-like behavior, in the present study, the results in the Vogel conflict test were in line with the contextual and trace fear conditioning tests, but not with the open field and elevated plus maze tests. One potential reason for this difference could be that positive allosteric modulation of GABA_A_ receptors by diazepam (the “baseline” in the Engin et al. 2016 study), which in all tested mice, including the conditional α2 knockouts in DG, CA3, and CA1 principal neurons, still acted at α1-, α3-, and α5-GABA_A_Rs and thus a potentially altered the excitation/inhibition balance altering computational integration of vCA1 inputs which results in a shift of the response to the approach-avoidance conflict of the VCT from anxiety-like to fear-related circuitry. It is noteworthy that both in the previous and in the current study the baseline number of licks, i.e., without shocks before the test session was similar in all the groups. An activation of NMDA receptors has been shown to be involved in the anxiety-like behavior displayed by rats in the VCT(30). While the previous study focusing on diazepam-induced inhibition of DG, CA3 and CA1 principal neurons identified a segregation between fear-related (CA1) and anxiety-like (DG, CA3) behaviors, the present study identified the roles of two major inputs to vCA1 connectivity with respect to fear-related (LEC-CA1, vCA3-vCA1) and anxiety-like (vCA3-vCA1) behaviors, independently of changes in the excitation/inhibition balance in other regions of the hippocampus.

The mechanisms regulating fear-related and anxiety-like behaviors in LEC to vCA1 projections are apparently distinct from that of vCA3 to vCA1 projections. While dorsal CA1 is innervated densely by medial EC layer 3 and sparsely by lateral EC layer 3(31), a recent study found that both dorsal CA1 and vCA1 receive stronger inputs from lateral EC rather than from medial EC(32). We also found that the projections from entorhinal cortex (layer 3) to CA1 contact in the stratum lacunosum-moleculare, whereas the projections from CA3 to CA1 contact in the stratum radiatum(24,25). However, in an anatomical study, it was found that all pyramidal neurons examined and 81% of the interneurons examined receive contacts from both projections(33). This study indicates that the target neurons may be largely the same for both projections, but that they target different layers of CA1 and thus their effects on the target neurons may be distinct. As identified in our previous work on conditional knockouts of the GABA_A_ receptor α2 subunit in CA1, CA3 and DG principal neurons, the inhibition of CA1 is necessary for diazepam-induced suppression of fear responses.

In the present study, which investigated the role of LEC projections to vCA1, neither activation nor inhibition of EC projections to vCA1 alter anxiety-like behavior in any of the anxiety-related assays including the elevated plus maze test, novel open field test, and Vogel conflict test. Together with the results discussed earlier, these findings indicate that the vCA3 to vCA1 projections modulate anxiety-like behavior but not the lateral EC to vCA1 projections. Our findings demonstrate that specific projections within hippocampal circuits play distinct roles in regulating anxiety-like behaviors, while they may be similar roles in regulating fear-related behaviors. This suggests that at a given time or a given anxiogenic context, of the two projections to vCA1, the vCA3 to vCA1projection may be actively engaged in modulating the computation in vCA1 such that the output is directed to the downstream brain regions that are involved in generating anxiety-like behaviors (19, 21).

In contrast to the functional divergence in these two hippocampal pathways with respect to anxiety-like behavior, both pathways did modulate fear expression behavior. The activation or inhibition of either the vCA3 projections or LEC projections to vCA1 led to an increase or decrease in contextual fear expression, respectively. These results are in line with another study reporting that vCA3 to vCA1 (vCA3-vCA1) projections promote contextual fear responses(34). In addition, previous lesion studies showed that vCA3 is important for retrieval of contextual fear conditioning and the vCA1 is important for retention of trace fear conditioning(35,36). The role of LEC in fear-related behaviors is also supported by other studies reporting that the activation of vCA1 neurons projecting to amygdala (18) or activation of an EC projection to the amygdala are required for contextual fear (18) or trace fear memory expression(37). is A potential limitation in our findings regarding fear-related behaviors in the LEC–vCA1 pathway is that, we observed that only one direction of optogenetic manipulation produced an effect: i.e., optogenetic inhibition of LEC projections in vCA1 leads to reduction of the freezing behavior in the tone test, whereas activation of the same projections did not alter freezing. However, in the context test, bidirectional optogenetic modulation was observed. Also, we noted that the freezing levels are different in sham controls of the two projections (i.e., vCA3–vCA1 and LEC–vCA1 projections). and the significance of this is unclear. The experiments were not performed on the same time or day for these projections.

Although both vCA3 and entorhinal cortical projections converge in vCA1 and collectively influence its activity, they exhibit distinct characteristics in their regulation of fear and anxiety. While both the temporoammonic pathway from the LEC to vCA1 and the trisynaptic pathway from the EC via vDG and vCA3 to vCA1 originate in the entorhinal cortex, previous studies suggest they process information differently. These pathways also originate from different layers within the EC: layer II neurons project to the DG, whereas layer III neurons project directly to CA1. Additionally, vCA3 receives afferent connections from several sources apart from dentate gyrus, including the entorhinal cortex, medial septum, and recurrent CA3 loops(25). Structurally, vCA3 neurons possess longer dendrites than dorsal CA3 (dCA3) neurons and display a higher ratio of recurrent CA3 inputs relative to dentate gyrus inputs when compared to dCA3 neurons(38). This suggests that vCA3 neuronal activity may be predominantly influenced by non-dentate afferents. This could potentially contribute to the differential effects of two projections that synapse on to vCA1.

Additionally, our findings and those of other research groups have demonstrated that CA3 and entorhinal cortex (EC) projections synapse at distinct layers within CA1, potentially contributing to the differences in information processing. In support of this, Vago et al reported that EC-to-CA1 projections are modulated by dopamine, a mechanism not shared by CA3-to-CA1 projections(39). Moreover, evidence from Brun et al revealed that while the direct entorhinal-CA1 pathway supports the acquisition of spatial recognition memory, effective spatial recall requires CA3-CA1 connectivity (40). Collectively, these studies highlight that CA3 and LEC inputs to vCA1 carry distinct information and separate modulations, likely explaining their differing roles in modulating fear-related and anxiety-like behaviors. To our knowledge, our study is the first to investigate the EC projections to vCA1 in the context of fear and anxiety-like behaviors. Further investigation of the cell type-specific responses in individual projections and their modulation by other neurotransmitters is warranted to further understand the difference between the two projections to vCA1.

In summary, our findings demonstrate a dissociation within the ventral hippocampus with regards to fear-related and anxiety-like behaviors, where vCA3 principal projections to vCA1, but not EC projections to vCA1, modulate anxiety-like behavior, while both vCA3 projections to vCA1, and LEC projections to vCA1 play a role in modulating fear expression. Our study highlights that the connection between the CA3 and CA1 regions plays a critical role in regulating anxiety-like responses, distinguishing it from other pathways that are involved in emotional regulation, such as the temporoammonic hippocampal pathway. These findings suggest that targeting the vCA3 to vCA1 pathway could be useful for the development of novel treatments for anxiety disorders.

## 4. METHODS AND MATERIALS

### 4.1 Animal subject

Procedures were conducted in accordance with the Guide for the Care and Use of Laboratory Animals of the National Research Council of the National Academies, and with approval from the University of Illinois Institutional Animal Care and Use Committee. One month old female and male C57BL/6J mice (stock #000664) were supplied by the Jackson Laboratory, Bar Harbor, ME. Both male and female mice were used for experiments starting at 10 weeks of age and provided with unrestricted access to food and water. Mice were housed 5 per cage on a 12 h reverse light/dark schedule with lights off at 11:00 a.m. and lights on at 11:00 p.m., with ambient temperature 23 ± 1 °C, and 50 ± 5% humidity. Experiments were conducted during the dark phase. Experiments for each of the pathways (vCA3 to vCA1, and lateral entorhinal cortex (LEC) to vCA1 optogenetic studies including activation (n=10), inhibition (n=10), and sham control (n=10) were performed with a total of 30 mice (15 males and 15 females). Equal numbers of male and female mice were used. 3 mice from the vCA3 to vCA1 projection group, and 6 mice from the LEC to vCA1 projection group have been excluded from the results due to off-target fiber optic ferrules or headstage detachment prior to study completion.

### 4.2 Viral constructs

Adeno-associated viruses for optogenetic manipulations were packaged and supplied by the UNC Vector Core Facility, NC. The vectors are as follows: AAV-CaMKIIa-EYFP (3.6×1012 mg/ml) opto-control, AAV-CaMKIIa-eNpHR3.0-EYFP (5.2×1012 mg/ml) opto-inhibition; AAV-CaMKIIa-hChR2(H134R)-EYFP (4.1×1012 mg/ml) opto-excitation.

### 4.3 Stereotactic surgeries

For all surgical procedures, mice were anesthetized with isoflurane (2–3%) and maintained under anesthesia (1-2%, 1 l/min O2) throughout the surgery and head-fixed in a stereotaxic frame (David Kopf, Tujunga, CA). Ophthalmic ointment was applied for eye lubrication, fur was shaved, the incision site sterilized, and body temperature maintained with a heating pad. Subcutaneous saline and ketoprofen (5 mg/kg) were provided preoperatively and postoperatively for up to 3 additional days for maintain hydration and analgesia, respectively as necessary.

For bilateral optogenetic illumination experiments, undiluted opsin viral vector was injected at the following coordinates in different animals (60-100nl volume per DV site); ventral CA3: −3.15 AP, 3.0 ML, −3.5 DV, lateral EC: −4.0 AP, 3.25 ML, −4.8 DV) and the needle was left in place for an additional 8-10 min to permit diffusion. Fiber optic ferrules were implanted bilaterally at the coordinates ventral CA1: −2.7 AP, 3.4 ML, −3.5 DV. After surgical procedures, mice were kept warm on a heating pad until they recovered. Mouse chow moistened with water was placed in the cage to initiate eating and provide hydration. Mice were allowed three weeks of recovery before starting any experiments.

Bilateral optogenetic manipulations. Three weeks after viral vector injection and fiber-optic implantation, mice were handled and habituated to fiber-optic adapter cables for 2 days prior to beginning behavioral experiments. For AAV-CaMKIIa-hChR2(H134R)-EYFP illumination, 5 ms pulses of blue light generated by a 50 mW 473 nm DPSS laser (OEM Laser Systems, UT) were delivered bilaterally via an optical fiber at 10-15 mW power presented at 20 Hz, and laser output was manipulated with an optic shutter controller (Thorlabs, Newton, NJ). For AAV-CaMKIIa-eNpHR3.0-EYFP illumination, constant yellow light illumination was generated by a 100 mW 593.5 nm DPSS Laser (OEM Laser Systems, Draper, UT) and delivered bilaterally via an optical fiber at 10 mW power. For all illuminations, the laser was split equally by a 1×2 Fiber Optic Rotary Joint (Doric Lenses, Canada) and delivered bilaterally through two implanted optical fibers connected to the optic patch cords using ceramic or metal sleeves (Thorlabs, Newton, NJ). A video camera and EthoVision XT15 software (Noldus, Leesburg, VA) were used to record live tracking of mice during behavioral experiments.

### 4.4 Behavioral experiments

All fiberoptic ferrule-implanted mice were subjected to behavioral experiments. All animals underwent blue or yellow laser illumination during behavioral experiments. Half of the control or sham group animals are subjected to illumination by either blue or yellow lasers. All the behavioral experiments including assessment of anxiety (Elevated plus maze, novel open field, and Vogel conflict test) and trace fear conditioning (contextual and tone fear testing), were performed similar to Kambali et al(20). For more details, follow the supplementary file 1.

### 4.5 Immunofluorescence and c-fos staining

Perfusion and preparation: Mice were deeply anesthetized with Ketamine / Xylazine (139 mg/kg / 21 mg/kg) and were perfused transcardially with ice-cold phosphate-buffered saline (PBS), followed by ice-cold 4% paraformaldehyde. The brains were dissected and post-fixed in the same fixation solution for 24 hr and transferred to 30% sucrose solution for cryoprotection for about three days. The brains were sectioned coronally into 40 mm-thick sections using a cryotome and collected as floating sections in PBS. These sections were divided for immunofluorescence, and for c-fos staining. Further details are provided in the Supplementary file 1.

### 4.6 Statistical analysis

For in vivo optogenetic illumination-coupled behavioral tests, statistical analysis was performed using Prism 9 (GraphPad Software, MA). Results were analyzed using unpaired Student t-test or one-way ANOVA with Sidak’s multiple comparisons post-hoc tests. Results are represented as mean ± S.E.M. P values <0.05 were considered significant. For multiple comparisons, p-values were adjusted accordingly.

## DATA AVAILABILITY

Supplementary figure 1, extended methods, and materials are provided in supplementary file 1. All data presented in this manuscript are available in supplementary file 2.

## ACKNOWLEDGEMENTS

This study was supported by grant #2017-08-31 from the Whitehall Foundation to UR.

## AUTHOR CONTRIBUTION STATEMENT

Maltesh Kambali (Investigation, Methodology, Formal Analysis, Data Curation, Writing—Original Draft, Writing—Review & Editing), Muxiao Wang (Formal Analysis, Writing—Review & Editing), Rajasekar Nagarajan (Formal Analysis, Methodology, Writing—Review & Editing), Jinrui Lyu (Formal Analysis, Writing—Review & Editing), Howard Gritton (Formal Analysis, Writing—Original Draft, Writing—Review & Editing) and Uwe Rudolph (Conceptualization, Formal Analysis, Methodology, Project Administration, Supervision, Funding Acquisition, Writing—Original Draft, Writing—Review & Editing)

